# How to stand out to a mate - Flash colour of *Photinus* fireflies maximizes signal contrast against ambient light

**DOI:** 10.64898/2026.10.29.760408

**Authors:** Judson Stricklin, Roland Kimsey, Simran Shah, Mrunal Nazare, James Masters, Kathrin F. Stanger-Hall

## Abstract

During twilight, flying male *Photinus* fireflies flash in search for conspecific females, which sit on the vegetation below and flash back in response. However, before darkness sets in, the green vegetation in the habitat makes the detection of flashes more difficult. Twilight active fireflies tend to emit more yellow flashes than nocturnal fireflies and it has been proposed that firefly flash colour evolved to enhance the contrast of firefly flashes against their signalling background. There is indirect evidence for this contrast hypothesis, but it remains to be rigorously tested. This is the first study to directly test the contrast hypothesis by recording ambient light spectra in three different habitats throughout the daily flash activity of *Photinus pyralis* and *P. australis* and comparing these ambient light spectra to the male flash spectra of the two species. The contrast hypothesis predicts that during peak activity, firefly light colour will maximally contrast with the signalling background. Our data show that male flash spectra fit into a gap in the ambient light spectrum that opens during peak flashing activity of each species, effectively enhancing the visual contrast between male flashes and their background and making males more visible to their potential mates.

## Introduction

Animal signals evolve to convey information between senders and receivers in their specific environments. As a result, signal production, the characteristics of the habitat through which the signal is transmitted, and the sensory properties of the receivers drive signal evolution. The sensory drive hypothesis [1] predicts that signals are under selection to maximize signal-to-noise ratio for the targeted receiver and are thus more easily detected in each environment, while detection by non-target receivers should be more difficult [2,3]. One mechanism to facilitate signal detection is to make signals more conspicuous, i.e. to increase signal contrast against the background.

For visual signals, such a high signal contrast has been documented for many animal groups, including insects [4], fish [2,5-8], lizards [9-11] and birds [12]. They all display colours that contrast with the environmental backgrounds against which they display. Many species of fireflies (Coleoptera, Lampyridae) also use visual light displays to attract and locate conspecific mates [13-15]. In many North American fireflies, flying males display short flashes that are repeated in species-specific flash patterns during mate search, while looking for response flashes from sedentary females [13-14]. The timing of these flash displays is key for species recognition and predator avoidance [16]. For example, character displacement in male flash duration suggests that flash duration is under selection in *Photinus* fireflies to reduce mating mistakes between sympatric species, and predation pressure by sympatric *Photuris* species seems to be selecting for extended flash pattern intervals, i.e. longer dark periods between repeated *Photinus* flash patterns [16]. Firefly species also differ in the colour (i.e. wavelengths with peak intensity) of their emitted light [17-19], however, only flash patterns, but not light colour, have been implicated as a factor in mate choice [20-21].

To explain the observed 30 nm range of firefly light colour, which - as perceived by human eyes-ranges from green (550 nm) to yellow-orange (579 nm), Lall and coworkers hypothesized that firefly light colour evolved to increase the contrast with the signal background and thus facilitates signal detection by potential mates [18]. This signal background can drastically differ between firefly species. Some species, especially those in the genus *Photinus*, start flashing well before sunset and continue during twilight. This includes the well-known Big Dipper (*Photinus pyralis*) fireflies that are widespread across Eastern North America and may signal in habitats ranging from forests and forest edges to open fields and roadsides [19,22]. Their activity extends into suburban yards, making them the most familiar firefly species in North America. The males of these slow-flying fireflies flash in regular intervals and hover at a height of ∼ 1m between flashes to wait for a female response flash from below, then fly ∼2-3m horizontally and repeat their display by dipping down and then flying back up during their flash, generating their characteristic J-shaped flashes. Other firefly species are active in the dark, including *Photuris, Pyractomena* and *Phausis* fireflies. The genus *Photuris* includes predatory fireflies that hunt other fireflies, including *Photinus*, and it is assumed that *Photuris* predation has facilitated the activity shift of *Photinus* fireflies towards earlier times in the evening, or even become diurnal, to escape predation [23].

This activity shift comes at a cost. Fireflies active at night experience a static (dark) signalling background without the interference of ambient light, except when a full moon is visible [24-25]. Light pollution contains different wavelengths at different intensities across the spectrum, depending on the anthropogenic light source [25-27], which may interfere with the production and/or detection of firefly light signals in different ways [28-32]. In contrast, early active fireflies experience a dynamically changing ambient light environment with a shifting distribution of light intensities across different wavelengths as the overall light level decreases. Approaching sunset, the ambient light is a mix of long (orange-red) wavelengths from the sky and shorter (green) wavelengths reflected off the vegetation [24-25]. After sunset, ambient light generally contains a higher proportion of short (blue) wavelengths, and as it gets dark, the ambient light spectrum is shifted back towards longer wavelengths (e.g. from star light, without moon) or is spectrally neutral (i.e. at full moon; [25]). As a result, the signal background of twilight-active firefly species continuously changes as the sun is setting, making it challenging for these species to produce light signals that stand out against the background enough to be detectable by conspecific mates.

The peak sensitivity of most insect eyes is in the green range [33-35] and this was likely also true for the first bioluminescent fireflies, which seem to have produced green light emissions [36]. Importantly, there is a close match between signal colour and visual sensitivity in fireflies [33] and other bioluminescent beetles [37]. The green flashes of dark-active *Photuris versicolor* and *Photuris lucicrescens* fireflies are matched by a green peak in their visual sensitivities [38-39] and the visual sensitivity of these fireflies reaches its maximum at the onset of darkness and lasts throughout the night [40]. Similarly, the green-yellow flashes of *Photinus pyralis* and the yellow-orange flashes of *P. scintillans* fireflies are associated with matching green-yellow and yellow-orange peaks in their visual sensitivity [33]. The contrast hypothesis explains this yellow-shift in both flash colour and visual sensitivity of early active fireflies as an adaptation to their ambient light environment [18], resulting in an increased signal-to-noise ratio against the green vegetation. This was generally supported by a survey of 32 dark-active species (72% emitted green light with peak emissions < 559 nm) and 23 dusk-active species (91% emitted yellow light with peak emissions > 560 nm; [18]) and by a modelling analysis [41-42]. However, these analyses did not account for differences in signalling habitat or the shared evolutionary history of the firefly species. A subsequent phylogeny-based analysis with habitat information and population-averaged light spectra for 23 firefly species, provided further support that earlier firefly activity is associated with yellow-shifted light emissions, but their results also modified the contrast hypothesis to include habitat type [19]. Specifically, early populations in more closed habitats (i.e. in forests, surrounded by green vegetation) emit flashes that are significantly more yellow-shifted compared to the flashes of early populations active in more open habitats (i.e. above fields or meadows), and thus were driving this trend [19]. These results further highlighted the importance of the different visual backgrounds for the evolution of flash colour, and the logical next step is to directly measure the ambient light in different habitats, while fireflies are signalling. This is the purpose of the present study.

We measured ambient light spectra for two twilight active *Photinus* species in three different habitats and recorded male flash spectra in each habitat. Based on the contrast hypothesis, we predicted that the male flash spectra of a given firefly species will maximally contrast with their ambient light environment at the peak of their flashing activity, when most mating signals are produced. We further predicted that fireflies that signal surrounded by vegetation and in ambient light environments that are richer in green wavelengths should have a more yellow-shifted flash colour than fireflies signalling in open environments.

## Methods

### I. Data collection

We collected flash count data, ambient light spectra and firefly flash spectra for two early active firefly species, *Photinus pyralis* (Big Dipper fireflies) and *P. australis* (Twilight Bushbabies) in three habitats during June and July 2024. For each species, flash counts and ambient light data were collected from the beginning of firefly activity (first observed flash) to the “end” of activity when flash counts had dropped to 10-30% of peak flash activity.

#### Field Sites

Two of the three field sites (Hilltop, Riverbottom) were located within Whitehall Forest and 0.85 km (straight-line distance) from each other, the third field site (Oakbend) was in the suburban Oakbend neighbourhood (∼ 9.5 km from the other sites). The Hilltop site (∼30m x 150m) was located in hardwood forest with an open understory (i.e. closed habitat of Hall et al. 2016), the Riverbottom site consisted of a mowed meadow (∼ 15m x 100m), separated by tree lines from two rivers that came together at this site, and a steep hillside forest on the other side (mixed habitat of Hall et al. 2016). The Oakbend site was in a more open suburban habitat with patches of hardwood trees, interrupted by large lawns and big houses. The Oakbend site consisted of a large lawn area (∼ 40m x 80m), separated by tall, dense Chinese Holly bushes and individual trees from roads on two sides of the property, and a hillside with hardwood trees and bamboo on the third side; the house was located on the fourth.

#### Firefly Species and Flash behaviour

The two twilight-active *Photinus* species occurred together in all field sites. *P. pyralis* males tend to be larger (9-19 mm) than the males of *P. australis* (8-11mm), and the two species can be distinguished by their different flash patterns [16,22]: (1) the yellow-green flashes of *P. pyralis* males are longer (0.6-0.8 seconds at 22-26°C) than the yellow-orange and “snappy” flashes of *P. australis* males (0.1-0.2 seconds at 22-26°C); (2) male *P. pyralis* emit a flash every 5-7 seconds (at 22-26°C), while *P. australis* flash every 1.5-4 s (at 23-27°C). In addition, the long *P. pyralis* flashes are emitted during an upwards swoop during flight (“J” shape), after which the male hovers in place to look for a female response below, before moving horizontally and flashing again about 2-3 meters away, while *P. australis* males fly in a short hopping motion right above the forest floor. Like *P. pyralis* males, *P. australis* males hover briefly after they flash, then fly a short distance before they repeat their flash [22]. In contrast to the low-flying (∼0.5m) *P. australis* males, *P. pyralis* males typically fly and signal at a height of ∼1m early in the evening and fly higher as it gets darker before retreating up into the trees as their display ends. The sedentary females of both species tend to respond from low vegetation to a conspecific male flash by aiming their light organ (on their ventral abdomen) towards the male and flash a single response flash with their species-specific flash delay and flash duration [14,16,22].

#### Flash Counts

We used flash counts for each species to determine the start of firefly activity and the time when most flashes were observed at a given field site (i.e. the peak activity time for each species). To monitor more of the habitat at each field site, the firefly habitat was divided into 5 or 6 individual observation areas, whose left and right boundaries were marked with orange flags and wooden stakes. Observers were sitting in camping chairs and counted all flashes they could see between their lateral markers in their line of sight. Each of the observers used a 30 second timer with an auditory end signal for regular flash counts (every 2-3 minutes) and recorded each flash count, along with the time of their count. The observers used flash duration (and visual up-swoops during flashes) to distinguish the two species. The flashes for each species were recorded in separate and alternating 30 second intervals. During peak activity times hand-held clickers were used to record flash counts.

#### Ambient Light spectra

Ambient light spectra were recorded using a pair of stacked Jaz spectrophotometers with two Oceanoptics UV-VIS fibers (300-1100nm). These were placed in one of the individual observer areas. To sample across habitat microsites at each field site, the ambient light spectrophotometers were placed within two different areas across the nights at each site. The light sensor of one of the Jaz spectrophotometers was positioned at 2-3m height, facing upwards towards the sky, (sky sensor: view of sedentary females as they look upwards towards flashing males), the sensor of the second spectrophotometer was positioned at ∼1m height, facing down to the ground (ground sensor: view of airborne male fireflies as they look for female response flashes below). Spectral readings were recorded every two minutes as photon counts and/or irradiance, resulting in 30 ambient light spectra each hour (30 sky spectra and 30 ground spectra) and saved as .txt files on a dedicated laptop computer, using the Oceanoptics software. To account for different integration times across days and sites, we translated all data into photon counts per 0.1 seconds before analysis (Table S1).

#### Firefly flash spectra

We used a dedicated Jaz spectrophotometer and Oceanoptics QP 600-2VIS-BX fiber (300-1100nm) to record flash spectra of individual firefly males during one of the last days of observation at each field site. All specimens with flash data were preserved as flash vouchers.

#### Environmental data

Ambient light levels influence the onset of firefly activity [43] and these light levels are influenced by time of sunset and cloud cover [24-25]. Air temperature affects firefly flash duration and flash repeats and thus flash counts [14] and drought may influence firefly survival and population size. To track the potential correlation of these and other environmental variables with firefly flash behaviour across observation days, we retrieved environmental data from NOAA (National Oceanic and Atmospheric Administration), which were collected at the National Weather Service station located within 10.5km (straight-line distance) from all three field sites (Table S2). In our figures we indicate cloud cover with the symbols used by NOAA [44] and moon phase with symbols of “timeanddate” [45].

### II. Data Analysis

#### Firefly activity

For each field site and evening, we used the flash counts for each species to determine their respective peak activity. Data from individual observation sites within a field site were analysed together to represent the firefly activity across the habitat. When multiple observers recorded a flash count for the same minute, their observations were averaged (arithmetic mean) for that time point. We report the time of highest flashing activity (max N flashes per 30 seconds), but rather than relying on a single data point (and possible outlier), we used the smoothed spline curve of the flash count distribution to generate an activity curve for the evening (JMP v18 graph builder, with a default lambda=0.05, trim=0.00 and confidence fit for the distribution spline) to determine the overall peak activity for each species. The confidence band for the spline fit in Jmp is empirically estimated using repeated bootstraps (resampling of data with replacement) to construct a confidence band that contains 95% of the smoother fits at each point along the curve (2025 JMP® v18. JMP Statistical Discovery LLC, Cary, NC). We used the Spline peak times to select the associated ambient light spectra for further analysis.

Flash counts over 30 seconds cannot be directly translated into firefly population counts since the males of *Photinus* fireflies signal in flash patterns with flash repeats every 5-7s for *P. pyralis* and every 2-4s for *P. australis* at environmental temperatures ranging between 22° and 26°C [22]. If a given male repeats his flash pattern for the whole 30s, about ∼5 flashes would be counted per *P. pyralis* male and ∼10 flashes per *P. australis* male, depending on the temperature. However, males occasionally insert longer pauses into their flash pattern series [22] and they may enter or leave the observation area during the 30s flash count, reducing the expected flash counts per individual. But given their similar male mate search behaviours and their different flash intervals, we would expect about 2 times as many *P. australis* flashes as *P. pyralis* flashes during the 30s counts at similar population numbers.

#### Ambient light spectra

For spectral analysis, the photon count files were loaded into R using the “getspec ()” function of the pavo module [46]. The recordings from the downwards facing (ground) fiber were too dark (ambient light levels too low) for a spectral analysis, therefore we focused our analysis on the ambient light spectra recorded by the upwards pointed (sky) light sensor, corresponding to the female view of male flash signals and their ambient light background. For each habitat we analysed the ambient light spectra recorded from 60 (HT) or 30 (RB, OB) minutes before sunset to 20 minutes after sunset (in 20- or 10-minute intervals) to document changes in ambient light throughout firefly activity. To directly compare the wavelength distribution in the ambient light spectra across time, we imported the raw ambient light spectra into Microsoft Excel (using the write.xlsx function) and normalized each spectrum to remove differences in overall brightness (luminance) in the ambient light as the sun sets. To normalize the spectra, we used a min-max normalization of photon counts for each data point with λ’ indicating the normalized data point, λrepresents the unnormalized data point, λ_*max*_ is the maximum value in the light spectrum, and λ_*min*_ the minimum value:

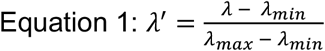

#### Contrast Hypothesis

To test the hypothesis that firefly flash colour in early active fireflies evolved to maximize flash contrast with the ambient light background, we predicted that in the dynamically changing ambient light environment that is experienced by early active fireflies such as *P. pyralis* and *P. australis*, the contrast between their flashes and ambient light should be highest during their peak activity when most flashes are produced as mating signals. To test this, we compared the normalized ambient light spectrum recorded at peak flashing activity (peak of spline in flash counts) for each species with the normalized ambient light spectrum recorded 30 minutes before peak activity (i.e. before or right after start of flashing activity). To visualize this wavelength shift, we calculated the difference in intensity (Δλ’) for each wavelength between these two normalized ambient light spectra:

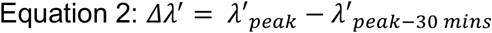

The absence of a colour (i.e. wavelength distribution) difference between these two ambient light spectra would result in a straight line across wavelengths (Null hypothesis). We then compared any wavelength shift in ambient light with an average flash spectrum recorded for the respective species at the same site.

#### Firefly light spectra

Firefly light spectra were analysed with a custom R script. Following [19], we fitted a cubic polynomial to the raw measurements to determine the peak wavelengths. For each firefly, we calculated the mean wavelength across its individual recorded light spectra, as well as the standard deviation and range. For each field site we averaged the individual means to calculate the mean peak wavelength (and standard error) for each species at that site. To ensure that only high-quality recordings of flash spectra were included in mean and standard deviation calculations, we filtered out measurements with a saturated amplitude (light intensity), as their peak emission wavelengths could not be determined accurately. In addition, spectra with a very weak signal (<5000 photon counts at peak wavelength), or disturbances, i.e. a peak value far outside the typical flash peak range (<525 nm or > 580 nm) for fireflies were also removed. To compare mean male flashes at the three sites we used a Games-Howell test (for unequal sample sizes and sample sizes 6+) after testing for equal variances.

#### Contrast as seen by fireflies

Animal signals and their contrast with the environment should be analysed based on the sensory system of the receivers, therefore firefly visual sensitivity must be considered [24,47]. The visual sensitivity of *P. pyralis* eyes has two peaks, one in the UV range (400 nm) and one in the long-wavelength LW (565 nm) range [33]. This is mediated by two different opsins, a UV opsin and a LW opsin [48-49]. Since firefly flashes do not contain a UV component and *Photinus* females only respond to wavelengths above 480 nm, only LW opsins are implicated in the achromatic detection of firefly mating signals [48,50]. Among fireflies, LW opsins, combined with eye screening pigments, provide a visual sensitivity spectrum that closely matches the flash spectrum for each species [33,38-39,51]. Therefore, we can use the flash spectrum, which represents not only the signal generated by the signalling male, but also the (slightly more narrow) visual sensitivity spectrum of conspecific receivers, with screening pigments excluding some of the greener and redder wavelengths [33:Fig 4B].

### Results

Both *P. pyralis* and *P. australis* were present in all three habitats, but due to an extended drought during this study their population numbers diminished over time (Fig. S1).

#### Flash spectra

We recorded the flash spectra of individual male *P. pyralis* from all three sites, and male *P. australis* flash spectra from the Hilltop site. Since the male *P. australis* were not flashing after being caught, they were taken to the lab and recorded inside clear 50ml Falcon tubes (do not impact light spectra) on the following evening, outside at ambient temperatures and ambient light. With 3-5 *P. australis* fireflies in each tube, they were able to move freely, increasing the likelihood of flashing. We report data based on a total of 5 high-quality light spectra for the whole group of *P. australis* from the Hilltop site (N= 10 individuals; Table S5), rather than from individual fireflies. We also recorded flash spectra from a single unidentified *Photuris* firefly from the Riverbottom habitat.

In the Hilltop habitat, the males of the two *Photinus* species flashed in the open understory of the hardwood forest, with brown leaves covering the forest floor amid sparse low growing vegetation. The two species flashed with significantly different flash colours (*P. pyralis* 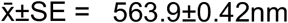, *P. australis* 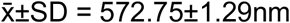; Wilcoxon Z=3.033, p=0.0012; Fig. 1A). In the suburban Oakbend habitat the fireflies were active over the large lawn area, along bushes and under a few tall trees at the edge of the property. The two roads bordering the property had only light neighbourhood traffic (∼ 5 cars per hour). A dusk-activated streetlight near the intersection of the two roads turned on during firefly activity. Most of the observation area was shielded from the light by tall (5-6m) dense Chinese Holly bushes, but a few *P. pyralis* fireflies were observed flying on the side of the bushes facing the streetlight. Compared to the more yellow flashes in the Hilltop habitat (hardwood forest: 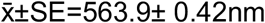), *P. pyralis* flashes in the Oakbend habitat were significantly greener (suburban yard: 561.79±0.43nm; Fig. 1B, Games Howell, q=2.68, p=0.0117). In the Riverbottom habitat, the fireflies were active above a mowed meadow, separated by tall grass and a few trees from two rivers, and bordered by tall grass and a steep hillside forest on the third side. In this habitat firefly activity started under the trees at the hillside edge of the meadow and moved into the open area. *P. pyralis* flashes in this mixed habitat 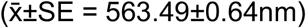 spanned the colour range of both the Hilltop and Oakbend habitats (Fig. 1B). In the Riverbottom habitat we also recorded a *Photuris* species, which was active at the end of *Photinus* activity and flashed with significantly greener flashes 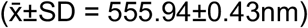 in this habitat than *P. pyralis* (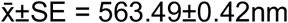; Wilcoxon Z= -3.36, p=0.0004; Fig. 1A).

**Figure 1:**
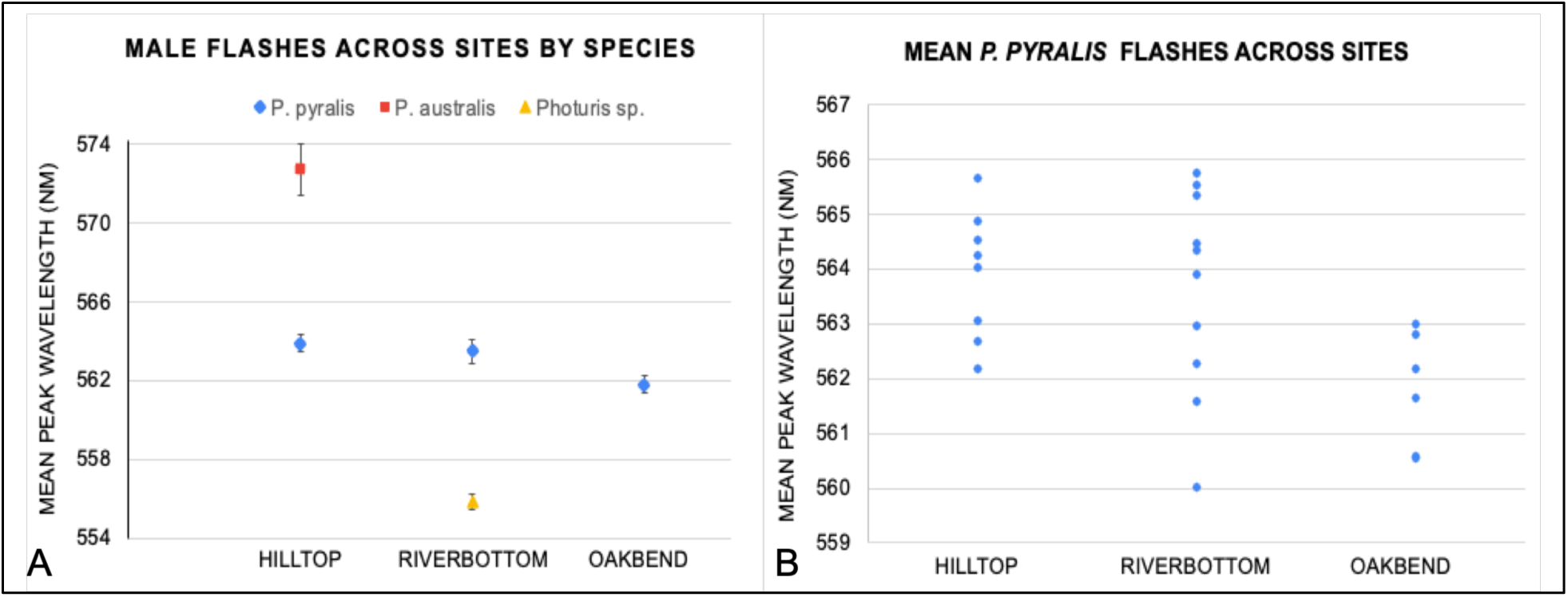
Male flash colour (mean peak wavelength of flashes) across sites. **A**. Blue diamond: mean 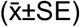 peak wavelength for *P. pyralis* flashes recorded at each site (based on individual means -HT: N=8, RB: N=10, OB: N=6); red square: mean 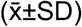 peak wavelength for male *P. australis* flashes recorded at the HT site; yellow triangle: mean 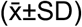 peak wavelength for the flashes of a *Photuris* sp. individual at the RB site. **B**. The distribution of mean individual male *P. pyralis* flashes across sites.

#### Flash activity across sites

Across habitats and environmental conditions (days since last rain, cloud cover, moon phase) the time of both the start and peak of firefly activity differed relative to sunset, but in each habitat both species were active approximately at the same time (Fig. 2). At the Hilltop site, both *P. pyralis* and *P. australis* were active at relatively high numbers. Both species started their activity earlier (100-110 min before sunset) on a day with overcast skies and a waxing crescent moon (June 12). Under clear skies the next day (June 13) they started 10-20 minutes later (∼90 min before sunset). The latest onset of flashing (∼60 min before sunset) occurred on days that were mostly clear with a waxing gibbous moon (June 18 and 20; Table S3). Over the four observation days at the Hilltop, *P. pyralis* started flashing 55-100 minutes before sunset and *P. australis* started 65-110 min before sunset (Table S3) in the same habitat. *P. pyralis* reached the peak of their flash distribution 18-46 minutes before sunset and *P. australis* 11-48 minutes before sunset (Fig. 2A, Table S3). Both species showed greatly diminished activity by sunset on all days at this site (Fig. 2A).

**Figure 2:**
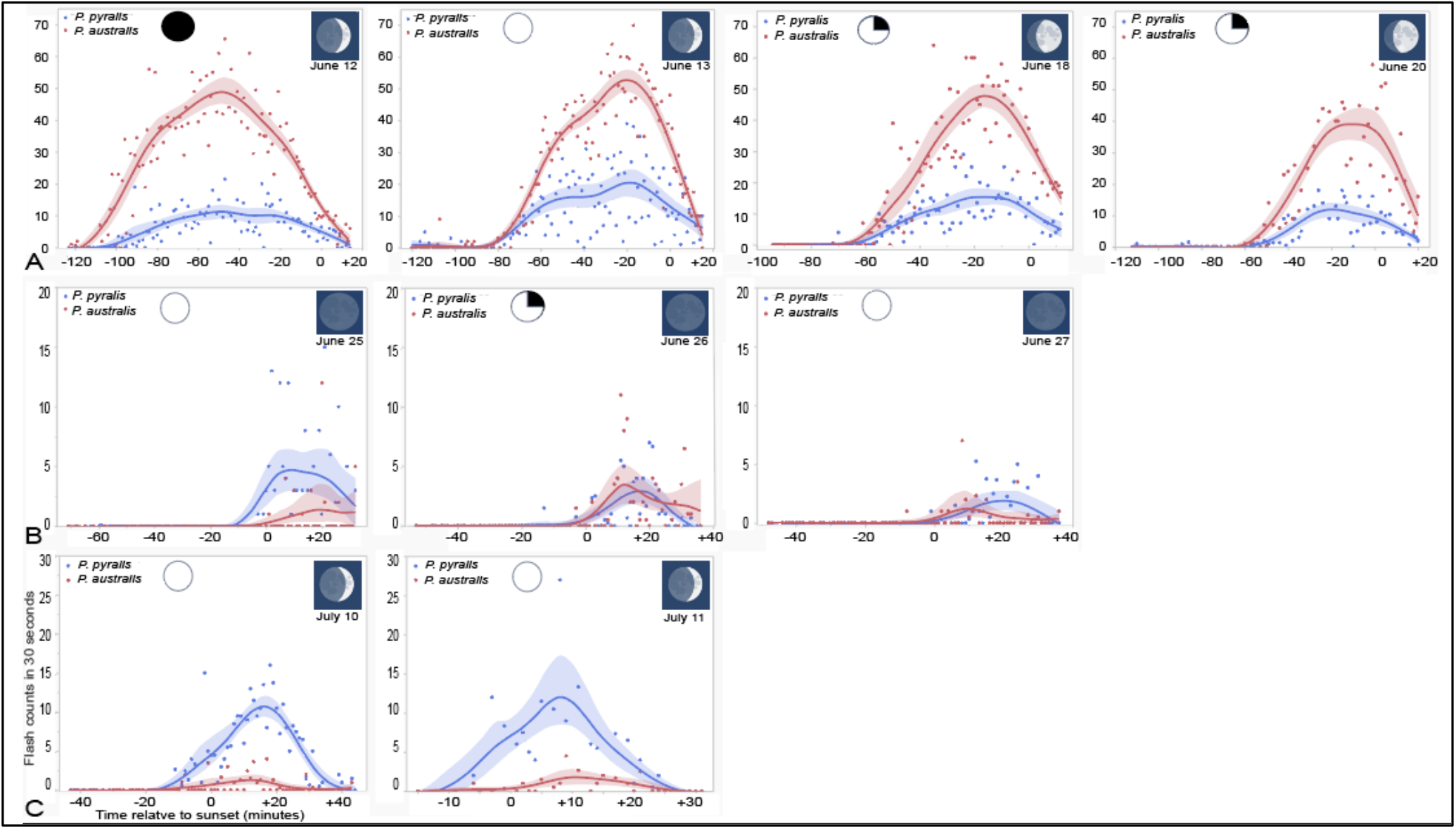
Flashing activity of *P. pyralis* (blue) and *P. australis* (red) in minutes before (-min) and after (+min) sunset (0min) at the A: Hilltop, B: Riverbottom, and C: Oakbend sites by date. Each data point represents the average flash counts (per 30s) for *P. pyralis* or *P. australis* fireflies across the individual observation areas (microhabitats). The peak of the spline curve (shown with confidence of fit) was used to determine the time of peak flashing activity for each species on each day and to select the corresponding ambient light spectra for analysis. Circle (black portion): cloud cover; square: moon phase (see also Table S2).

In the Riverbottom habitat (2 weeks after the last rain: Fig. S1), both *Photinus* species were active at relatively low numbers from 5-10 min before sunset across the three observation days. *P. pyralis* reached peak activity 9-20 minutes after sunset and ceased activity ∼ 30 min after sunset (Fig. 2B). The few *P. australis* at this site reached peak activity 9-19 min after sunset and ceased activity soon after. A *Photuris* sp. firefly was observed flashing 30 minutes after sunset (at 21:18) on June 27 in this habitat.

In the suburban Oakbend habitat, *P. pyralis* and especially *P. australis* were active at low numbers, with *P. pyralis* numbers somewhat recovered after it had rained on June 27 (Fig. S1). *P. pyralis* started flashing 5-15 min before sunset, and *P. australis* 0-5 minutes before sunset, their flashing activities peaked 12-17 min after sunset, and both stopped flashing by 30-40 min after sunset (Fig. 2C, Table S5).

#### Ambient light across habitats

To track the changes in ambient light during firefly activity before and after sunset in each habitat, we analysed the ambient light spectra every 20 minutes, starting at 60 min before sunset until 20 min after sunset in the Hilltop habitat, and every 10 minutes, starting at 30 min before sunset until 20 min after sunset in the Riverbottom and Hilltop habitats. Different wavelengths dominated (highest photon counts) the ambient light spectra in the different habitats (Fig. 3): wavelengths ∼ 550 nm (green) dominated the ambient light at the Hilltop site (hardwood forest; Fig. 3A); at the Riverbottom (meadow between forest and river) and Oakbend (suburban yard) sites, the most dominant wavelengths were ∼ 510 nm (blue-green; Fig. 3B&C). During firefly activity, the overall light intensity (photon counts) diminished across sites, but the dominant wavelengths stayed the same until sunset.

**Figure 3:**
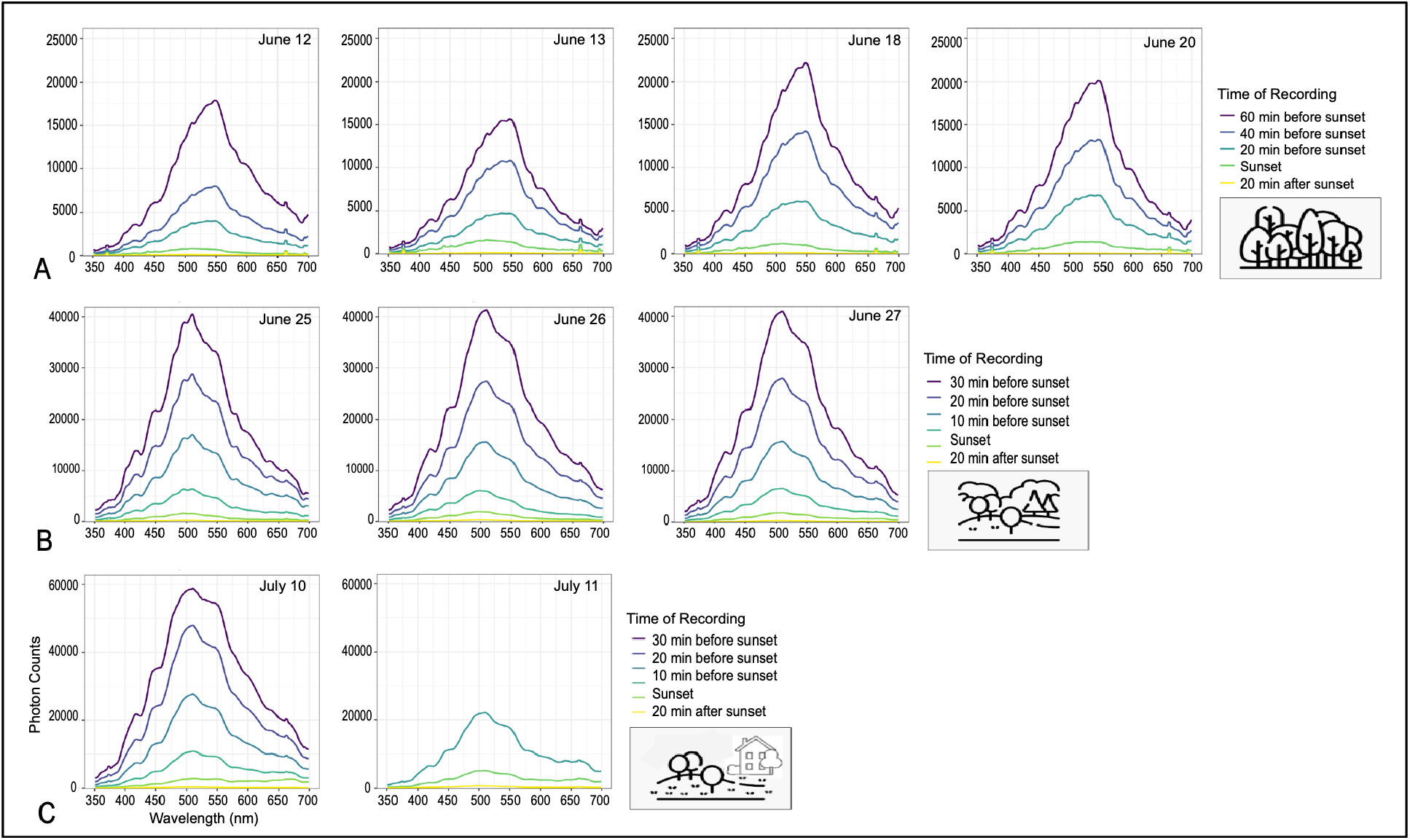
Ambient light spectra (photon counts across wavelengths) by site and date at A: Hilltop site (forest), B: Riverbottom (meadow), and C: Oakbend (suburban yard) sites, starting at 60 minutes (Hilltop) or 30 minutes (Riverbottom and Oakbend) before sunset until 20 minutes after sunset. For clarity, we show the ambient light spectra in 20- or 10-minute intervals. The artificial noise blips <375nm and >660nm were filtered out before normalizing the spectra. On July 11 ambient light recordings started within 10 minutes of sunset.

**Figure 4:**
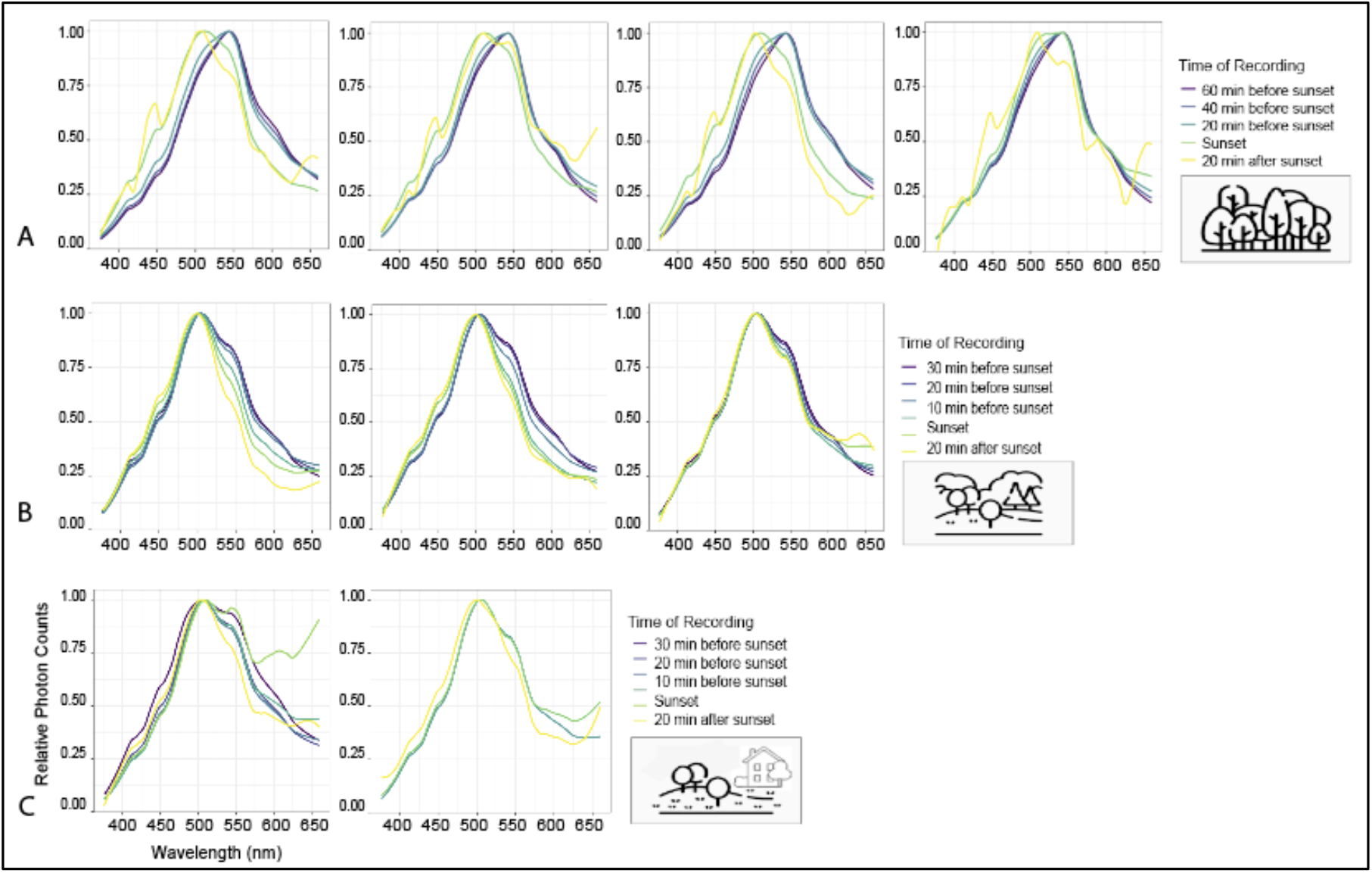
Normalized and smoothed ambient light spectra (after bandpass filtering) for different habitats and days in 20-(HT) or 10-minute (RB, OB) intervals from 60 or 30 minutes before to 20 minutes after sunset. A: Hilltop (forest), B: Riverbottom (meadow), C: Oakbend (suburban yard).

Some of our raw ambient light spectra showed consistent noise blips at the two ends of the spectrum (at 372-375 nm and at 661-664 nm: see Fig. 3), therefore we applied a bandpass filter to exclude all wavelengths < 375nm nm and > 660 nm in our ambient light spectra before normalizing the ambient light spectra for further analysis.

#### Ambient light and flashing activity

To separate light intensity from the wavelength distribution in ambient light we plotted the normalized ambient light spectra for each species at each site and observation day. Due to the high measurement noise in the normalized ambient light spectra recorded after sunset (small measurement fluctuations in these low intensity measurements are disproportionately amplified during normalization: Fig. S2), we smoothed the normalized spectra to recognize trends (Fig. 4). Across all habitats and days, the smoothed normalized ambient light spectra shifted towards shorter wavelengths as it got darker. The dominant wavelength shifted at sunset from ∼550 nm to ∼510 nm in the Hilltop forest and from ∼510 nm to ∼500 nm at Riverbottom and Oakbend (Fig. 4).

The hypothesis that firefly flash colour evolved to contrast with the signalling background (ambient light) at peak flashing activity predicts that there is less interference of ambient light with the flash spectra of *P. pyralis* (yellow-green flashes) and *P. australis* (yellow flashes) males when they are flashing. To test this, we plotted the wavelength shift curve (i.e. the difference in the wavelength distribution of the normalized smoothed ambient light spectra) that had occurred at peak flashing activity compared to 30 minutes earlier, when flashing had just started or was about to start, for each species at each site and observation day, and compared it with the average flash spectrum for *P. pyralis* or *P. australis* at the respective site.

In all instances, the ambient light spectra recorded at peak flashing activity of *P. pyralis* contained fewer wavelengths that would overlap with the flash spectrum of *P. pyralis* males compared to 30 minutes earlier (Fig. 5). The same is true for *P. australis*. Like *P. pyralis* flashes (Fig. 6A), the flashes of *P. australis* (Fig. 6B) closely match a gap in ambient light wavelengths that opens during their peak flashing activity at the Hilltop site (deciduous forest), despite their flashes being significantly shifted towards longer wavelengths (*P. australis* 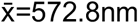 versus *P. pyralis* 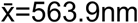). In the Riverbottom habitat, *Photuris* fireflies were observed flashing their greener flashes (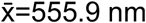 versus *P. pyralis* 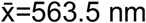) on June 27 at 30 minutes after sunset. At this time, the ambient light was at very low levels across wavelengths (i.e. too low to register, e.g. Fig. 3B) and the activity of both *Photinus* species had almost completely ceased (Fig 2B).

**Figure 5:**
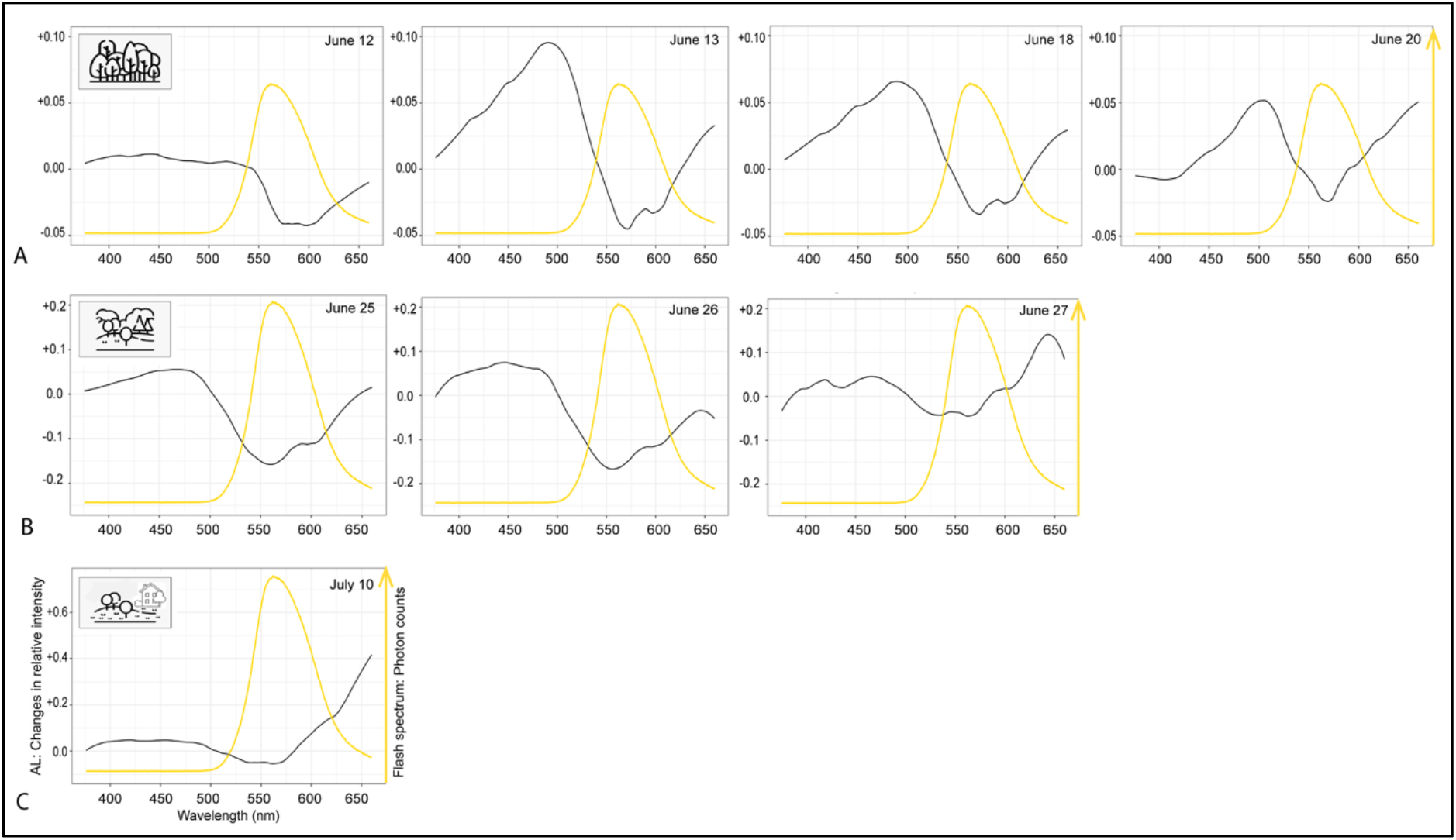
Average flash spectra of *P. pyralis* (yellow line: photon counts across wavelengths; not standardized) recorded at each site, superimposed on the wavelength shift in ambient light (black line: relative intensity difference across wavelengths) at the same site. A value of 0 shows that no change has occurred in the 30 minutes between start and peak of flashing activity. A: Hilltop (forest), B: Riverbottom (meadow), C: Oakbend (suburban yard; recordings on July 11 started after 30 min before peak and not included here).

**Figure 6:**
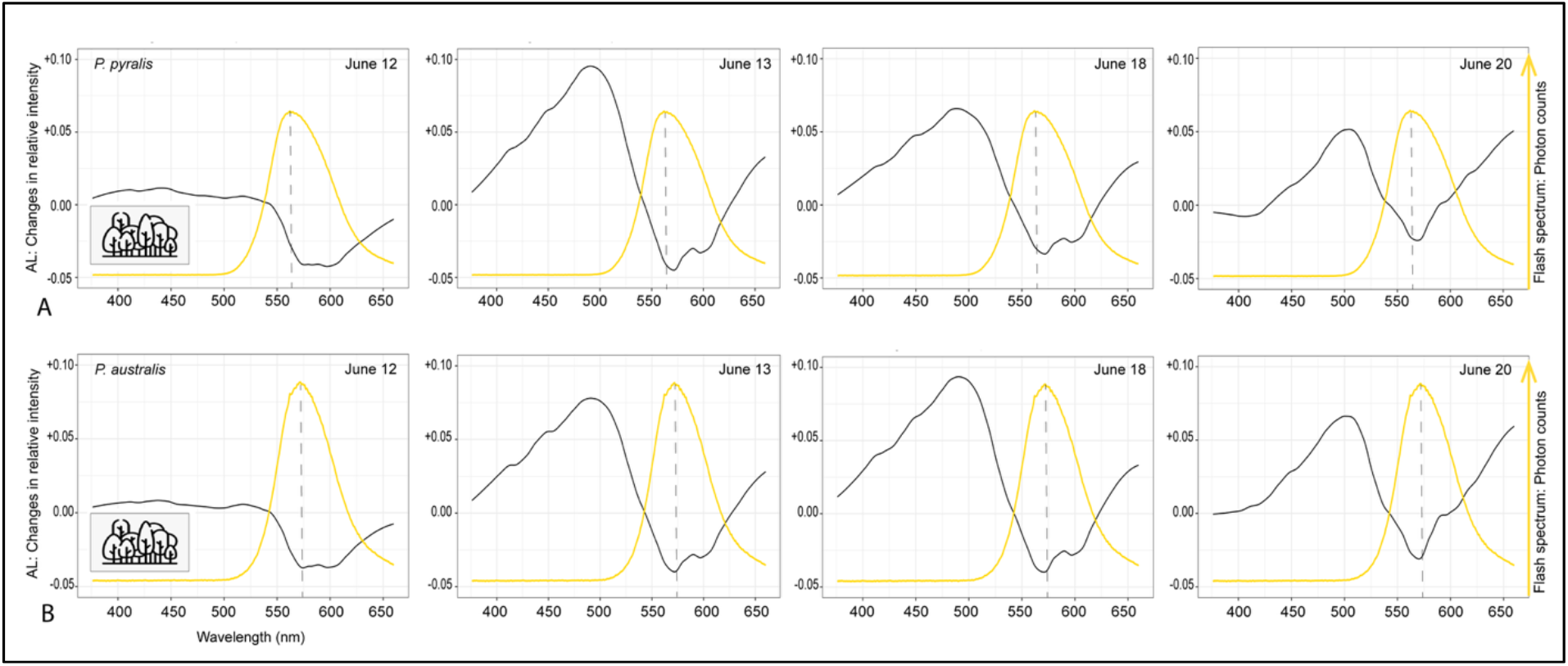
A: *P. pyralis* and B: *P. australis* average flash spectra (yellow curve: photon counts across wavelengths; not standardized) at the Hilltop forest site, superimposed on the ambient light wavelength shift (black line) at that site. A value of 0 shows no change has occurred in the 30 minutes between start of flashing and peak flashing activity. The stippled line marks the wavelength with peak intensity of the average flash of each species in the Hilltop habitat (deciduous forest).

## Discussion

Our ambient light spectra, combined with firefly flash spectra from the same habitat, provide unambiguous support for the contrast hypothesis, which explains the evolution of yellow-shifted flash colours in early active fireflies as an adaptation to stand out to potential mates against the still visible green background in their habitat [18]. Here we document the dynamically changing ambient light environment in three different habitats, with a shifting distribution of light intensities across different wavelengths as it gets darker. These are the conditions that *P. pyralis, P. australis*, and other early-active fireflies experience during their mate search. As noted by Endler [24] light in forests is unique. While the dominant wavelengths in the Hilltop forest were green (∼550 nm), the Riverbottom meadow and the Oakbend yard were dominated by blue-green light (∼510 nm) with the sky contributing more blue wavelengths. This wavelength dominance did not change until sun set in each habitat (Figs 3,4), when the dominant wavelengths also started to shift towards shorter wavelengths (HT to ∼510 nm; RB and OB to ∼500nm). As predicted by the contrast hypothesis, the greener ambient light in the Hilltop forest (closed habitat of [19]) was associated with significantly more yellow *P. pyralis* flashes compared to the greener flashes in the suburban (open) habitat. *P. pyralis* in the Riverbottom meadow (mixed habitat) flashed across the colour range of both the Hilltop (closed) and the Oakbend (open) habitats, likely due to the combination of hillside forest and open meadow at this site. The ability of *P. pyralis* fireflies to move between closed and more open parts of habitats at some sites may explain why their flash colours do not always significantly differ between habitats [19].

Our study shows that firefly flash colour is an adaptation to a wavelength gap in ambient light that opens during peak flashing activity in their specific habitat, facilitating conspecific signal detection and potential mating success. Individual fireflies flash with similar peak wavelengths (within ∼2-3nm), but the variation between individuals generates a distribution of peak wavelengths in a population [19]. This distribution allows selection to act on firefly flash colour and these selective forces are expected to differ in different habitats. Indeed, previous studies have shown a large range of population means across the geographic distribution of *P. pyralis* [19,52] and our results provide convincing evidence that (1) the selective forces on light colour of a given population act through contrast optimization with the ambient light environment at peak flashing activity in their specific habitat, and that (2) the peak activity times and the ambient light spectra differ between habitats. In the forest habitat the overall activity time of *P. pyralis* started much earlier and lasted longer than in the open or mixed habitats. This may have partly been influenced by the higher population numbers in early June, but a longer activity time in forested habitats has also been observed for *P. scintillans* fireflies [53].

As sunset approaches, the ambient light tends to be a mix of long (orange-red) wavelengths from the sky and shorter (green) wavelengths from ambient light that is reflected off the vegetation [24-25], and this was also the case in our three habitats (Fig. 3). After sunset ambient light contains a higher proportion of shorter wavelengths [25], and in our three firefly habitats this is mainly due to longer wavelengths in the ambient light spectrum dropping out disproportionately. As a result the right slope of the ambient light spectrum is left-shifted towards shorter wavelengths (Fig. 4). This left shift opens a wavelength gap in ambient light (Figs. 5,6), which the firefly flashes of *P. pyralis* and *P. australis* have evolved to utilize for mating success, through both, flash colour and timing of flashing

### Flash colour of different firefly species

Differences in firefly flash colour, e.g. the significant difference between the flashes of *P. pyralis* (yellow-green) and *P. australis* (yellow) or *Photuris* (green) are easily distinguished by human eyes, however, they cannot be distinguished as different colours (chromatic vision) by fireflies [20,50]. Beetles have lost the ancestral blue-sensitive insect opsin [54]. Different groups of beetles have regained a third pigment by gene duplications of the UV-sensitive and/or LW opsin genes, however, this is not the case for fireflies, possibly influenced by their mostly dim environments, which are often associated with low opsin diversity in insects (e.g. Lepidoptera: [55]). As a result, fireflies have only two opsins, one UV and one LW opsin [48-49]. Two different flash wavelength distributions within the sensitivity range of one LW opsin, like the flash spectra of *P. pyralis* and *P. australis* cannot be distinguished as a different “colour” (chromatic difference) by fireflies but are detected achromatically instead [50]. For a chromatic distinction of the two *Photinus* species, two or more neurally integrated LW opsins with partially overlapping sensitivities would be needed [56]. The inhibition of mate attraction to green light by male *Lampyris* fireflies by adding blue light was interpreted as evidence for color vision [57], but the higher input to the UV/blue photoreceptor, typical for early evenings, may simply inhibit the activity of this nocturnal firefly. However, fireflies still can respond differently to light stimuli of different “colours” (wavelength distributions) if they are presented in the same ambient light background and one stimulus has a higher contrast with this background than the other. This higher signal contrast is a direct result of the interaction between the signal spectrum, firefly visual sensitivity and the composition of ambient light at that time.

### Tuning of flash spectra and visual sensitivity

For the tuning of firefly flash spectra to closely match their visual sensitivity, both signal detection (eyes) and signal production (light organs) are under selection [41-42]. The visual sensitivity of *P. pyralis* has two peaks, one in the UV (400 nm) and one in the long-wavelength LW (565 nm) range [33]. Since firefly flashes do not contain a UV component and *Photinus* females only respond to wavelengths above 480 nm, only LW opsins are implicated in the achromatic detection of firefly mating signals [50,19]. Screening pigments in the eyes of fireflies play an important role in tuning the visual sensitivity to the flashes of each species by filtering out certain wavelengths before the light hits the photosensitive cells [58], and screening pigments in light organs also seem to be responsible for individual and population-level flash variation [59]. Early active *Photinus* fireflies, like *P. pyralis* use pink screening pigments, which selectively filter out greener (and redder) wavelengths from the incoming light, making their visual sensitivity narrower, while shifting peak sensitivity into the yellow range [58]. In comparison, *Photuris* fireflies use yellow pigments (filtering out blue wavelengths) and day-active non-bioluminescent fireflies like *Lucidota atra* lack screening pigments altogether [41-42,51,58]. Opsin photoproteins do not seem to play a major role in visual tuning as shown by both, a modelling analysis [41] and a phylogeny-based molecular analysis of UV and LW opsins in 38 firefly species [48]. In firefly light organs, screening pigments can also be used to filter the light generated by luciferase, the enzyme catalysing the bioluminescence reaction, before it is emitted. A recent study documented 46 products in the ommochrome and pterin pigment pathways that are expressed in both the eyes and light organs of *P. pyralis* fireflies [59]. While mainly serving detoxification and tissue protection, these two pathways produce several coloured pigments and pigment precursors that could filter the emitted light in the light organ, as well as the light admitted to the photoreceptors in the eyes of fireflies. This is the strongest argument yet for the importance of pigment filters in both light organs and eyes, especially since flash colour differences between populations of *P. pyralis* are not due to amino acid differences in luciferase [52]. An analysis of light organ screening pigments across *Photinus* and *Photuris* fireflies is presently underway and will be able to shed more light on how the pigment filters may differ between species, and to what degree they may contribute to the tuning between signal production and signal detection across firefly species (and genera) in different light environments.

Despite all their benefits, pigment filters come at a cost by reducing signal brightness [51] and this cost is higher (i.e. brightness more reduced) the more yellow-shifted the signal is [42]. This seems counterintuitive, because signal brightness is important for signal contrast and reproductive success, therefore it is important to keep in mind that this reduction in brightness is disproportionately due to a reduction in the shorter (interfering) wavelengths, i.e. a marked attenuation in the green range (480-530nm) for *Photinus* fireflies [58]. This increases the relative visibility of the longer (more yellow or orange) wavelengths in the signal, even though the overall brightness is reduced. This is not unique for firefly mating signals [6]. Despite the reduction in overall signal brightness by screening pigments in the eyes and light organs of early-active fireflies, there are several alternative ways to enhance signal detection. This includes (a) relatively larger eyes in bioluminescent fireflies [60], (b) a larger light organ surface area (e.g. via a larger body size [61-62],) to generate a brighter signal, (c) a stronger reflector (dorsal uric acid layer) behind the firefly photic cells to reduce light scattering and emit a brighter signal [63], and (d) more oxygen admitted to the light reaction by the tracheal system to make the signal brighter [64]. In addition, the bioluminescence reaction could possibly be tweaked by (e) making more ATP and/or (f) luciferin (the substrate of the light reaction) for the light reaction [65], however, this also requires more luciferase to increase reaction turnover and to limit reaction inhibition. How much of the bioluminescent reaction chemistry is already optimized in the light organs of fireflies and which tweaks their homeostasis and energy balance would still allow, remains to be studied.

Once a male flash signal is detected, firefly females recognize a conspecific male by his flash duration [20,14,16], then point their light organ at the male and flash their response flash at a species-specific delay to the male flash and a flash dialogue ensues [20,14). All this is mediated by the LW opsin and visual screening pigments. The sensitivity of the UV opsin likely serves firefly navigation through the environment during UV-rich twilight and can also sense some blue light (400-500nm) that is abundant at that time [51]. Interestingly, UV and LW opsins are also expressed in firefly light organs [52], but their function, if any, remains to be determined. The only other known organism with opsins in a light organ is the squid *Euprymna scolopes*, who possesses the molecular, biochemical, and physiological capability for light detection in its light organ and presumably uses it to monitor its light-producing bacterial symbionts [66]. Why fireflies, who produce their own bioluminescence have opsins in their light organs and whether they can use them to sense ambient light levels to trigger their activity, or whether they are used to regulate luciferin synthesis in preparation for signalling remains to be tested.

### Timing of Flashing activity

The peak flashing activity of *P. pyralis* fireflies coincides with the opening of a wavelength gap in ambient light. This timing, combined with their yellow-green flashes that match this wavelength gap (Fig. 5), optimizes the chances of a signalling *P. pyralis* male to be detected by a female on the vegetation below and to elicit a female response. Our data (from the upwards facing “sky” sensor) represent this female perspective. From the male perspective our recordings from the downwards facing (“ground”) sensor were too dark (i.e. ambient light levels too low) for a spectral analysis. This means that the ambient light in our three habitats is less likely to interfere with signals emitted by females sitting on vegetation close to the ground and aimed at males that are looking down. In addition, *P. pyralis* females can emit significantly greener signals than *P. pyralis* males, which suggests that any green leaves from which a female signals, can act as reflectors, amplifying the greener female flashes [19] and making them more detectable by males.

Given the tight match between firefly flash spectra and the ambient light gap at peak flashing activity across all three habitats, why do some *P. pyralis* males flash so much earlier (up to 60 minutes at HT and up to 30 min earlier at RB and OB), when it is brighter overall, their signal contrast is lower and their flashes therefore much more difficult to detect by females? The benefits of flashing earlier include the avoidance of flash competition with many other males (followed by scramble competition for mating when a female responds: [67]) and the high reproductive pay-off when a female does respond. *P. pyralis* females can mate multiple times, but only once per night [68]. This means that any mated female will be removed from the mating pool for the night, leading to declining female numbers throughout the activity time. So why do not more males start flashing early? The short answer is that male resources are limited. Except for some nectaring [69], non-predatory fireflies do not tend to eat as adults and all their resources for mate search stem from larval fat deposits. In addition, *P. pyralis* males transfer nuptial gifts during mating, which come at a cost [70]. Therefore, daily mate search time is negatively correlated with the number of nights a male can search. If a male does not find a female, he may continue searching past peak activity, when signal contrast increases due to low ambient light, however, this comes with a decreasing probability of still finding a female and an increasing risk of predation by *Photuris* fireflies. As a result, the exact timing of male flashing in a population is a trade-off between costs and benefits incurred by individual males (Fig. 7). We predict that based on the shifting wavelengths in ambient light, males with more yellow flashes in the population should be relatively more successful in being detected by females before peak activity, while males with greener flashes benefit from flashing around peak activity. This makes firefly flashing (timing and colour) a perfect example of sensory drive in action, with signal production, the characteristics of the habitat through which the signal is transmitted, and the sensory properties of the receivers driving signal evolution, resulting in a maximized signal-to-noise ratio for the targeted receiver, while detection by non-target receivers is more difficult [1].

**Figure 7:**
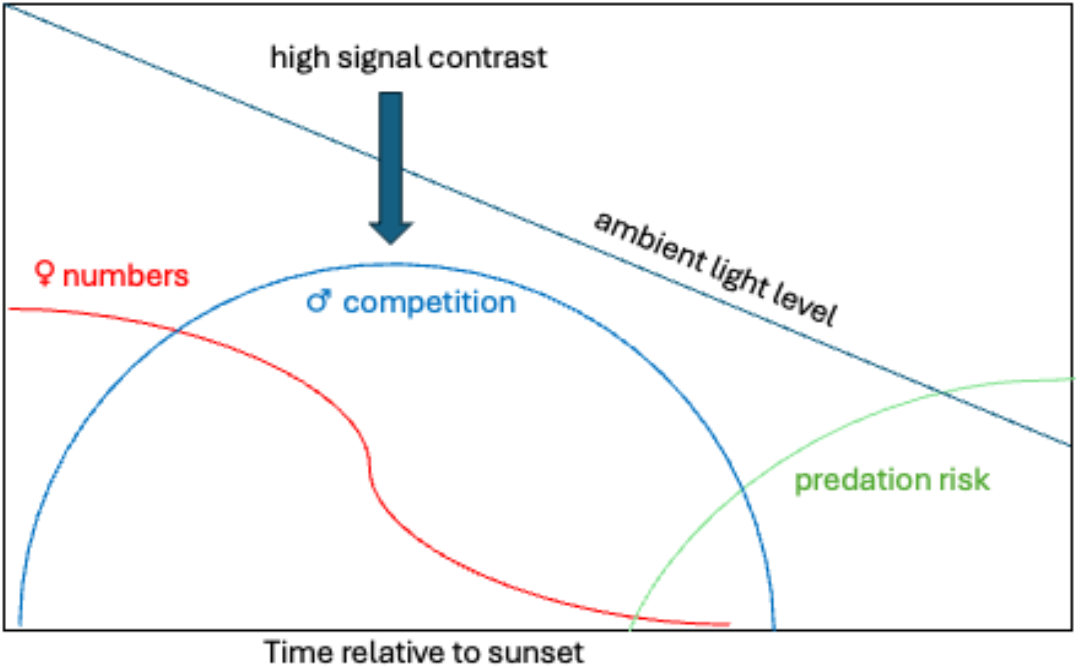
Costs and benefits of the timing of mate search (flashing activity) of individual firefly males. The numbers of unmated females decrease through the activity time, and predation risk increases as it gets darker. Male competition is highest at peak flashing time when signal contrast is higher than during earlier flashing.

### Environmental influences on firefly behaviour

Both firefly species in our study started their flashing activity at very different light levels (Supplement: III) across habitats and weather conditions (e.g. moon light and cloud cover). Fireflies use low ambient light levels to initiate their daily activity [43,29] and a good example of this was seen during the total solar eclipse in 2017 when fireflies started to flash from North Carolina to Missouri as the sun disappeared [71]. On the other hand, artificial light at night (ALAN) can have detrimental effects on firefly mate search behaviour [29, 32]. Light pollution, if present, contains different wavelengths at different intensities across the spectrum, depending on the anthropogenic light source [25-27], which may interfere with the production and/or detection of firefly light signals in unique ways [28-32], and females tend to mate only in dark areas [72]. The streetlight at the suburban Oakbend site came on during firefly activity. Even though we observed a few males actively flashing while exposed to the streetlight, *P. pyralis* females are less likely to respond under ALAN [72-74]. In suburban neighbourhoods, outdoor lights on houses are common. Considering that all LED light colours reduce firefly flashing activity to some degree [74], neighbourhoods can preserve their firefly populations by (a) switching lights on later in the evening, (b) putting all outdoor lights on motion sensors, and (c) install lights that point downwards, rather than spreading light pollution (ALAN) over a large area [75]. In addition to ALAN, insecticides targeting mosquitoes, low vegetation diversity, removal of leaf litter, municipal irrigation and fertilizers reduce firefly abundance in suburban yards and suggest opportunities for firefly conservation [76]. By protecting the mate search behaviour of bioluminescent fireflies, these efforts will also help conserve other beneficial flying insects that are attracted and possibly harmed by outdoor lights as well [77-78].

## Supporting information

Supplement Parts I-III

## Acknowledgements

We are grateful to the Warnell School of Forestry and Natural Resources at the University of Georgia for access to the Whitehall property and to the Lindenberg family for letting us use their yard as a research site. Thank you also to Desi Hoagland for helping develop the counting techniques, and to Mike Morris for technical support (Jaz spectrophotometers). Part of the equipment for this research was funded by a UGA seed grant to KSH, and JS was partially supported by a UGA CURO (Center for Undergraduate Research Opportunities) stipend.

