## Supplement Parts I-III for "How to stand out to a mate - Flash colour of *Photinus* fireflies maximizes signal contrast against ambient light"

#### Table of Content


**I. Supplemental Figures**
Figure S1: Days since last rain and peak flash activity across observation days.
Figure S2: Normalized ambient light spectra across sites.

**II. Supplemental Tables**
Table S1: Environmental Data
Table S2: Ambient light spectra integration times
Table S3: Flash activity of *P. pyralis* and *P. australis*: Date, field site and time of sunset.
Table S4: Flash spectra recorded from *P. pyralis*, *P. australis* and *Photuris* fireflies by field site
Table S5: Individual recorded light spectra

**III. Onset of daily firefly activity (lux measurements)**
Overview and Methods
Figure S3: Changes in lux for each habitat and day
Table S6: Light levels at start of activity (lux)

**IV. References for Supplement**

### I. Supplemental Figures

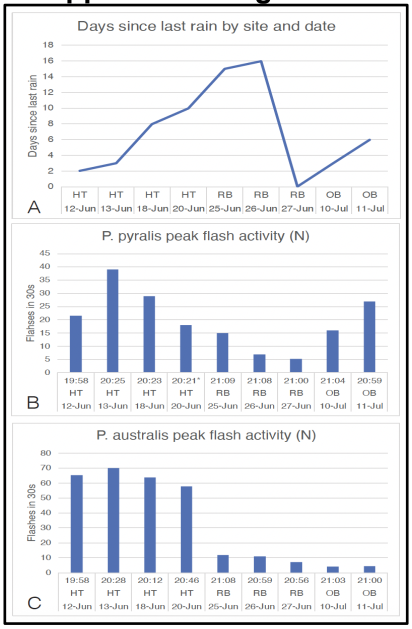


**Figure S1**: (A): Days since last rain and peak flash activity (max number of flashes in 30s) across days and field sites for (B) *P. pyralis* and (C) *P. australis*. The time at which peak flash counts (N) were recorded, the field site and the date are noted for each day. *A second peak (same flash number) was recorded at 20:32 for *P. pyralis on June 20.*

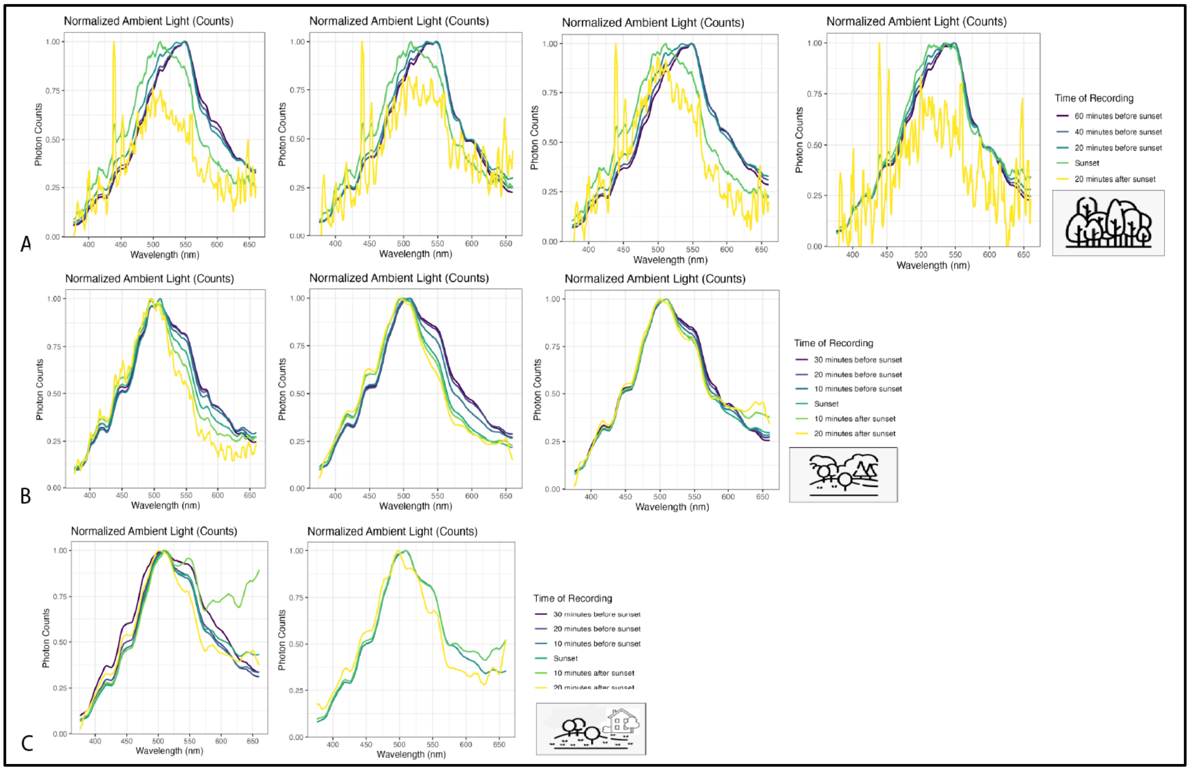


**Figure S2:** Normalized ambient light spectra for A: Hilltop (forest), B: Riverbottom (meadow) and C: Oakbend (suburban yard). During normalization small changes during low ambient light (i.e. 20 min after sunset) are disproportionately amplified, therefore we smoothed spectra before normalization for further analysis.

#### II. Supplemental Tables

**Table S1**: Environmental Data. All data were retrieved for 9pm if possible, otherwise for *8:00 PM; **firefly flash spectra recorded. Data retrieved from A: NOAA [1] and B: "timeanddate" [2] ^ the moon takes ~25 hours to rise and set, there was no moon rise on that day.

| Date (2024) | June 12 | June 13 | June 18 | June 20 | June 25 | June 26 | June 27 | July 10 | July 11 |
|---|---|---|---|---|---|---|---|---|---|
| Field site | HT | HT | HT | HT** | RB | RB** | RB** | OB | OB** |
| A: Temperature (°C) | 29.4 | 31.7 | 30.6 | 31.1 | 35.6 | 37.2 | 27.8 | 31.1 | 32.8 |
| A: Dew Point (°C) | 16.7 | 16.1 | 16.7 | 15.6 | 12.8 | 14.4 | 22.2 | 18.3 | 15.6 |
| A: Wind Speed (m/s) | 2.6 | 3.1 | 6.2 | 4.6 | 2.6 | 1.5 | 1.5 | 5.1* | 3.6 |
| A: Sky Cover (Oktas) | 8 | 0 | 2 | 2 | 0 | 2 | 0 | 0 | 0 |
| A: Sky Cover | Overcast | Clear | Mostly Clear | Mostly Clear | Clear | Mostly Clear | Clear | Clear | Clear |
| A: Last Rain | 6/10 | 6/10 | 6/10 | 6/10 | 6/10 | 6/10 | 6/27 | 7/9 | 7/9 |
| Days since last rain | 2 | 3 | 8 | 10 | 15 | 16 | 0 | 3 | 4 |
| A: Precipitation (mm) | 0 | 0 | 0 | 0 | 0 | 0 | 21 | 0 | 0 |
| B: Sunset | 20:45 | 20:45 | 20:47 | 20:47 | 20:48 | 20:48 | 20:48 | 20:46 | 20:46 |
| B: Moonrise | 12:06 | 13:03 | 17:49 | 19:58 | N/A^ | 0:05 | 0:37 | 10:52 | 11:48 |
| B: Moon Altitude | 52° | 54° | 28° | 9° | Below Horizon | Below Horizon | Below Horizon | 34° | 38° |
| B: Moon Heading | 238° | 220° | 149° | 133° | Below Horizon | Below Horizon | Below Horizon | 253° | 240° |
| B: Moon Phase | Waxing Crescent | Waxing Crescent | Waxing Gibbous | Waxing Gibbous | N/A | N/A | N/A | Waxing Crescent | Waxing Crescent |
| B: Moon Illumination | 38.2% | 47.9% | 90.7% | 97.0% | N/A | N/A | N/A | 22.7% | 31.40% |

**Table S2:** Integration times of ambient light measurements [3]

| Date | Integration time | Recorded as | Recorded as | Translated to |
|---|---|---|---|---|
| June 12 | 0.6 seconds | photon counts | irradiance | photon counts per 0.1 sec |
| June 13 | 0.3 seconds | photon counts | irradiance | photon counts per 0.1 sec |
| June 18 | 0.3 seconds | photon counts | irradiance | photon counts per 0.1 sec |
| June 20 | 0.3 seconds | photon counts | irradiance | photon counts per 0.1 sec |
| June 25 | 0.1 seconds | photon counts | irradiance | photon counts per 0.1 sec |
| June 26 | 0.1 seconds | | irradiance | photon counts per 0.1 sec |
| June 27 | 0.1 seconds | | irradiance | photon counts per 0.1 sec |
| July 10 | 0.1 seconds | | irradiance | photon counts per 0.1 sec |
| July 11 | 0.48 seconds | | irradiance | photon counts per 0.1 sec |

**Table S3**: Flash activity of *P. pyralis* and *P. australis*: Date, field site and time of Sunset. The time of the first flash and time of peak flash activity is listed for each species, date and habitat as clock time (24 hours) and in hours: minutes relative to sunset (Sunset=0, + after sunset, - before sunset). Start/Sunset: first flashes relative to sunset (Sset); Npeak/Sset: peak number of flashes (N/30sec) relative to sunset (hours: minutes); Speak/Sset: peak flashes (based on spline of flash distribution) relative to sunset (hours: minutes). We used the spline peak (Speak) to select the corresponding ambient light spectra for analysis.

| Location | | | *P. pyralis* | | | | | | | *P. australis* | | | | | | |
|---|---|---|---|---|---|---|---|---|---|---|---|---|---|---|---|---|
| Date | Site | Sunset | 1st Flash | Start/Sset | peak Time | Npeak (30s) | Start-peak | Npeak/Sset | Speak/Sset | 1st Flash | Start/Sset | peak Time | Npeak (30s) | Start-peak | Npeak/Sset | Speak/Sset |
| 12-Jun | HT | 20:45 | 19:04 | -1:41 | 19:58 | 21.5 | 1:48 | -0:47 | -0:46 | 18:43 | -2:02 | 19:58 | 65.5 | 1:15 | -0:47 | -0:48 |
| 13-Jun | HT | 20:45 | 18:47 | -1:58 | 20:25 | 39 | 1:41 | -0:20 | -0:18 | 18:58 | -1:47 | 20:28 | 70 | 1:30 | -0:17 | -0:20 |
| 18-Jun | HT | 20:47 | 19:37 | -1:10 | 20:23 | 29 | 0:46 | -0:24 | -0:21 | 19:35 | -1:12 | 20:12 | 64 | 0:37 | -0:35 | -0:16 |
| 20-Jun | HT | 20:47 | 18:57 | -1:50 | 20:21* | 18 | 1:26 | -0:26 | -0:19 | 19:47 | -1:00 | 20:46 | 58 | 0:59 | -0:01 | -0:11 |
| 25-Jun | RB | 20:48 | 20:45 | -0:03 | 21:09 | 15 | 0:24 | +0:21 | +0:09 | 20:49 | +0:01 | 21:08 | 12 | 0:19 | +0:20 | +0:19 |
| 26-Jun | RB | 20:48 | 20:35 | -0:13 | 21:08 | 7 | 0:24 | +0:20 | +0:17 | 20:45 | -0:03 | 20:59 | 11 | 0:14 | +0:11 | +0:12 |
| 27-Jun | RB | 20:48 | 20:47 | -0:01 | 21:00 | 5.25 | 0:13 | +0:12 | +0:20 | 20:40 | -0:08 | 20:56 | 7 | 0:16 | +0:08 | +0:09 |
| 10-Jul | OB | 20:46 | 20:35 | -0:11 | 21:04 | 16 | 0:29 | +0:18 | +0:17 | 20:37 | -0:09 | 21:03 | 4 | 0:26 | +0:17 | +0:12 |
| 11-Jul | OB | 20:46 | 20:45 | -0:01 | 20:59 | 27 | 0:14 | +0:13 | +0:13 | 20:45 | -0:01 | 21:00 | 4.5 | 0:15 | +0:14 | +0:16 |

**Table S4**: Flash spectra recorded from *P. pyralis*, *P. australis* and *Photuris* fireflies by field site and habitat (category of Hall et al. 2016). *P. australis**: A total of five high-quality spectra [5] were recorded from *P. australis* (N=10) as a group [10]; flash spectra were recorded from one *Photuris* specimen in the field. **SD: standard deviation from pooled specimens or individuals; SE: standard error of individual means.

| Site | Habitat (category) | Species | Individuals recorded (N) | N spectra/ individual | Mean peak wavelength (nm) | SD* or SE of means |
|---|---|---|---|---|---|---|
| Hilltop | wooded | *P. pyralis* | 6 | 3-21 | 563.84 | 0.38 |
| | | *P. australis** | [10] | [5] | 572.75 | 1.29** |
| Riverbottom | meadow | *P. pyralis* | 10 | 1-17 | 563.49 | 0.64 |
| | | *Photuris* sp. | 1 | 7 | 555.94 | 0.43** |
| Oakbend | suburban | *P. pyralis* | 6 | 1-6 | 561.79 | 0.43 |

**Table S5**: Individual firefly light spectra: mean, standard deviation (SD) and sample size (N) (numbering of individual spectra IDs restarts each day).

| Species | Date/Individual | Habitat | Recorded | Mean | SD | N | Individual Measurements |
|---|---|---|---|---|---|---|---|
| P. pyralis | Spectra062024/w01 | HT | field | 563.07 | 0.13 | 2 | 562.98, 563.16 |
| P. pyralis | Spectra062024/w02 | HT | field | 564.51 | 0.70 | 5 | 564.47, 564.36, 565.70, 563.92, 564.12 |
| P. pyralis | Spectra062024/w04 | HT | field | 564.02 | 1.35 | 7 | 564.68, 564.73, 563.83, 565.88, 563.04, 561.68, 564.28 |
| P. pyralis | Spectra062024/w11 | HT | field | 564.24 | 2.03 | 15 | 567.29, 565.63, 564.93, 568.90, 561.71, 562.94, 562.88, 563.34, 562.69, 565.72, 564.04, 562.39, 561.97, 564.34, 564.79 |
| P. pyralis | Spectra062024/w12 | HT | field | 562.67 | 0.38 | 3 | 562.24, 562.84, 562.93 |
| P. pyralis | Spectra062124/P1 | HT | next day | 562.17 | 1.15 | 9 | 561.07, 560.77, 562.21, 560.99, 561.69, 562.33, 563.59, 563.90, 562.97 |
| P. pyralis | Spectra062124/P6 | HT | next day | 564.88 | 0.67 | 5 | 565.34, 564.56, 565.10, 565.53, 563.88 |
| P. pyralis | Spectra062124/P11 | HT | next day | 565.67 | 1.57 | 6 | 566.22, 565.10, 565.56, 567.21, 562.93, 566.98 |
| P. pyralis | Spectra062524/P1 | RB | field | 564.47 | 1.72 | 9 | 562.07, 562.06, 564.55, 564.77, 564.92, 564.25, 564.79, 564.99, 567.84 |
| P. pyralis | Spectra062524/P2 | RB | field | 562.95 | 0.83 | 17 | 563.78, 562.67, 563.44, 562.73, 561.80, 563.42, 562.15, 564.24, 562.06, 564.00, 563.82, 563.95, 562.96, 562.59, 562.03, 562.68, 561.81 |
| P. pyralis | Spectra062524/P4 | RB | field | 563.90 | 0.78 | 4 | 562.85, 564.11, 563.92 564.73 |
| P. pyralis | Spectra062624/P2 | RB | field | 564.32 | 1.04 | 9 | 565.98, 564.42, 563.66, 564.20, 565.78, 564.08, 564.06, 562.55, 564.18 |
| P. pyralis | Spectra062624/P3 | RB | field | 565.51 | 1.64 | 3 | 564.12, 567.33, 565.10 |
| P. pyralis | Spectra062624/P4 | RB | field | 565.74 | NA | 1 | 565.74 |
| P. pyralis | Spectra062624/P5 | RB | field | 560.02 | 0.42 | 2 | 559.72, 560.32 |
| P. pyralis | Spectra062724/P1 | RB | field | 565.35 | 2.10 | 12 | 568.98, 561.49, 561.67, 566.62, 565.45, 567.21, 565.64, 566.02, 565.67, 564.72, 566.01, 564.72 |
| P. pyralis | Spectra062724/P2 | RB | field | 562.28 | 0.67 | 13 | 562.68, 562.05, 562.40 ,562.33, 562.48, 562.86, 562.55, 563.36, 562.11, 561.36, 561.41, 561.11, 562.99 |
| P. pyralis | Spectra062724/P3 | RB | field | 561.59 | 0.73 | 8 | 560.95, 562.27, 562.75, 560.84, 561.74, 561.06, 562.13, 560.97 |
| P. pyralis | Spectra071124/P1 | OB | field | 561.64 | 0.74 | 3 | 560.90, 561.65, 562.37 |
| P. pyralis | Spectra071124/P2 | OB | field | 562.98 | 1.30 | 4 | 561.82, 561.90, 564.24, 563.95 |
| P. pyralis | Spectra071124/P3 | OB | field | 562.16 | 0.85 | 6 | 561.11, 561.38 561.85, 562.51, 562.93, 563.19 |
| P. pyralis | Spectra071124/P4 | OB | field | 560.55 | NA | 1 | 560.55 |
| P. pyralis | Spectra071124/P5 | OB | field | 560.59 | NA | 1 | 560.59 |
| P. pyralis | Spectra071124/P6 | OB | field | 562.82 | 1.59 | 6 | 566.01, 561.91, 562.33, 562.38, 562.46, 561.80 |
| P. australis | Spectra062124/group | HT | next day | 572.75 | 1.30 | 5 | 571.52, 572.99, 573.99, 573.97, 571.28 |
| Photuris | Spectra062724 | RB | field | 555.85 | 0.40 | 10 | 556.87, 556.03, 555.63, 555.67, 555.77, 555.84, 555.78, 555.40, 555.91, 555.60 |

**III. Onset of daily firefly activity** (lux measurements)

The daily start of firefly activity is triggered by ambient light levels, which for *Photinus umbratus* average ~ 4 lux to start activity [4], but varies widely among fireflies [5]. Since UV and LW opsins may be involved in detecting the threshold of ambient light that cues the initiation of evening flashing activity [6,7] we used the whole light spectrum (390-700nm) in our spectra to calculate the light levels (lux) at which flashing started for each site and when it peaked for *P. pyralis* and *P. australis*. We used lux, a standardized measure of illuminance, which measures the amount of light falling on a surface per unit area. Lux is a standard measure for measuring brightness in the literature [5] but it weighs light wavelengths based on how human eyes perceive brightness, not necessarily how fireflies perceive it.

To calculate the light levels at the onset and the peak of activity for *P. pyralis* and *P. australis* in the three different habitats, we first transformed the recorded ambient light levels (recorded as irradiance) at these times from units of $\mu W/cm^2/nm$ to $W/m^2/nm$. After this conversion, these values were converted to Lux using the Equation 3 from [3] but with wavelengths limited up to 700nm (*P. pyralis* visual sensitivity) [4].

Equation 3: $Illuminance\ (lux) = k \int_{390}^{700} I(\lambda)E(\lambda)d\lambda$

In terms of habitat brightness (as perceived by human eyes), both species started flashing at 25-30 lux at the HT (forest) site and flashing activity peaked at 4-11 lux across the 4 observation days (with waxing crescent or waxing gibbous moon under overcast to clear conditions). At the RB (meadow) site both species started flashing at 2-5 lux and peaked between 0-1 lux across 3 observation days (with no moon visible and clear or mostly clear conditions). At the OB site (suburb) flashes started between 1-12.5 lux and peaked at 0.2-1 lux (with a waxing crescent moon and clear conditions; Fig. S3, Table S6)

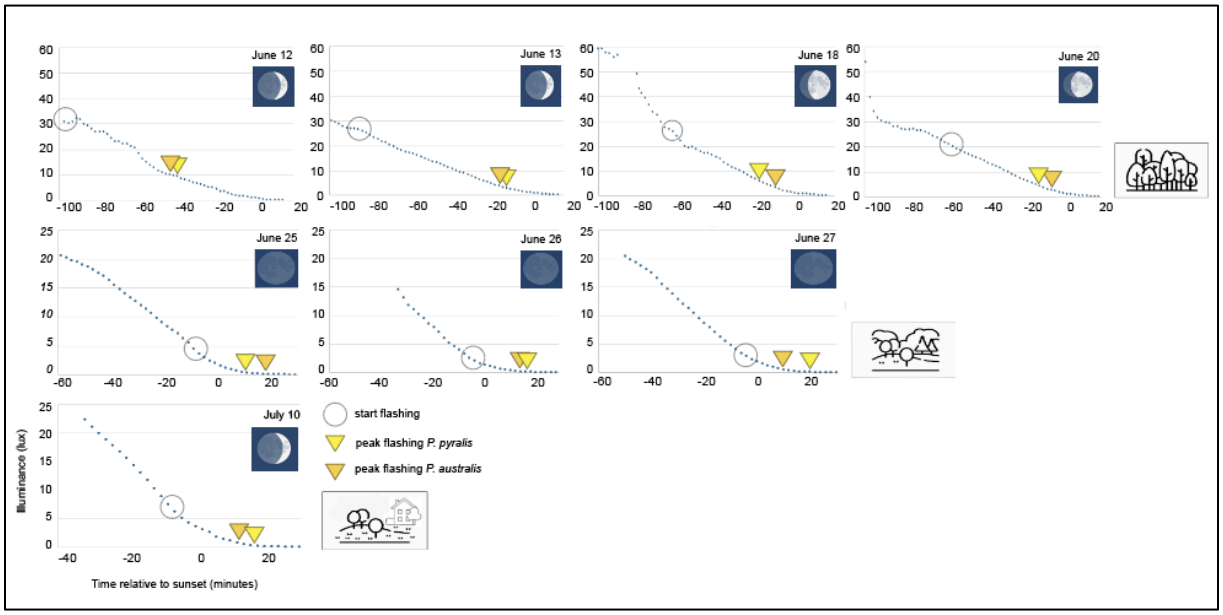


**Figure S3:** Illuminance (lux) changes over time across sites and days.

**Table S6**: Light levels (lux) at the start and at the spline peak (Speak) of firefly activity across habitats. Start and population peak times are given in minutes relative to sunset on each day (- before or + after sunset).

| Date | Site | *Photinus australis* | | | | *Photinus pyralis* | | | |
|---|---|---|---|---|---|---|---|---|---|
| | | Start-Sunset | Start (lux) | Speak-Sunset | Speak (lux) | Start-Sunset | Start (lux) | Speak-Sunset | Speak (lux) |
| June 12 | HT | -110 min | >30 | -48 min | 11 | -100 min | 30 | -46 min | 10 |
| June 13 | HT | -90 min | 27 | -20 min | 5 | -90 min | 27 | -18 min | 5 |
| June 18 | HT | -65 min | 30 | -16 min | 5 | -65 min | 30 | -21 min | 8 |
| June 20 | HT | -65 min | 25 | -11 min | 4 | -55 min | 30 | -19 min | 8 |
| June 25 | RB | -5 min | 2 | +19 min | 0 | -10 min | 5 | +9 min | 1 |
| June 26 | RB | -5 min | 2.5 | +12 min | 0.5 | -5 min | 2.5 | +17 min | 0 |
| June 28 | RB | -5 min | 3 | +9 min | 0.5 | -5 min | 3 | +20 min | 0 |
| July 10 | OB | -5 min | 5 | +12 min | 1 | -15 min | 12.5 | +17 min | 0 |
| July 11 | OB | 0 min | 1 | +16 min | 0.2 | -5 min | - | +13 min | 0.3 |

#### IV. References for Supplement


1. NOAA: https://www.weather.gov

2. TimeandDate: https://www.timeanddate.com/astronomy/moon/phases.html.
